# Background selection under evolving recombination rates

**DOI:** 10.1101/2021.12.20.473549

**Authors:** Tom R. Booker, Bret A. Payseur, Anna Tigano

## Abstract

Background selection (BGS), the effect that purifying selection exerts on sites linked to deleterious alleles, is expected to be ubiquitous across eukaryotic genomes. The effects of BGS reflect the interplay of the rates and fitness effects of deleterious mutations with recombination. A fundamental assumption of BGS models is that recombination rates are invariant over time. However, in some lineages recombination rates evolve rapidly, violating this central assumption. Here, we investigate how recombination rate evolution affects genetic variation under BGS. We show that recombination rate evolution modifies the effects of BGS in a manner similar to a localised change in the effective population size, potentially leading to an underestimation of the genome-wide effects of selection. Furthermore, we find evidence that recombination rate evolution in the ancestors of modern house mice may have impacted inferences of the genome-wide effects of selection in that species.

## Introduction

Different modes of selection (e.g. positive, purifying and balancing) all affect genetic variation at sites linked to the actual targets of selection (reviewed in Charlesworth 2009). In the case of purifying selection, the removal of deleterious mutations causes linked neutral variants to be lost along with them through a process referred to as background selection (BGS; Charlesworth et al. 1993). Of the mutations that affect fitness in natural populations, the vast majority are likely deleterious with a comparatively small proportion of beneficial mutations (Eyre-Walker and Keightley 2007). For those reasons, it has been proposed that BGS is ubiquitous across eukaryotic genomes and should be incorporated into null models for population genomics (Comeron 2017; Johri et al. 2020). Indeed, recent studies have used BGS to set baseline patterns for identifying the locations and effects of positively selected mutations (DeGiorgio et al. 2016; Campos et al. 2017) and understanding Lewontin’s paradox of genetic diversity (Buffalo 2021). Interpreting genome-wide patterns of genetic diversity in terms of BGS, however, requires accurate estimates of population genetic parameters, particularly recombination rates.

In many species, the recombination rate per base pair (*r*) varies across the genome both between and within chromosomes (Stapley et al. 2017). For example, in the house mouse (*Mus musculus*) the average *r* for chromosome 19 (the shortest chromosome) is around 60% higher than for chromosome 1 (the longest chromosome)(Cox et al. 2009). The requirement of at least one cross-over per chromosome per meiosis in mammals causes shorter chromosomes to recombine at a higher average rate than longer ones (Pardo-Manuel et al. 2001; Segura et al. 2013; Dumont 2017). Local recombination rates can vary substantially across chromosomes as well and in some cases this variation is predicted by gross features of chromosome architecture such as the locations of centromeres and telomeres (Paigen et al. 2008). Actual recombination events in mice are typically restricted to narrow windows of the genome (on the order of 1-5 Kbp), referred to as hotspots (Paigen et al. 2008). The positions of recombination hotspots in mice, and in some other vertebrates, are determined by the binding of a protein encoded by the *PRDM9* gene to specific DNA motifs (Baudat et al. 2010; Baker et al. 2017), although hotspots are still observed in *PRDM9* knockout lines and dogs, which lack a functional copy of *PRDM9* (Brick et al. 2012; Auton et al. 2013).

Estimates of *r* can be obtained empirically by examining the inheritance of genetic markers through controlled crosses or through pedigrees, or by comparing an individual’s genome to that of its gametes (e.g. Sun et al. 2019). Both methods reconstruct recombination events over one or a few generations, and thus provide estimates of *r* for contemporary populations. Alternatively, estimates of *r* can be obtained indirectly by analysing patterns of linkage disequilibrium across the genome (e.g. Spence and Song 2019), in which case estimates reflect both recent and ancestral recombination events. Whether recombination rates are estimated from marker transmission or population genetics, using such estimates when analysing of variation across the genome in terms of BGS implicitly assumes that the recombination landscape has not changed over the time in which patterns of diversity have been established. However, recombination rate landscapes can evolve very rapidly in some lineages. For example, due to the relationship between chromosome size and average *r*, changes in chromosome length (i.e. karyotype evolution) may induce changes in *r*. The lineage leading to *Mus musculus* (2*n*=40) has experienced large chromosomal rearrangements since it shared a common ancestor with *Mus pahari* (2*n*=48) 3-5 million years ago (Thybert et al. 2018). Moreover, different populations of *Mus musculus domesticus* harbouring different karyotypes exhibit different genomic landscapes of recombination (Vara et al. 2021). Chromosomal fusions can exhibit meiotic drive (Chmátal et al. 2014) so new karyotypes may spread to fixation very rapidly. Even mice with the same karyotype vary in regional recombination rate across substantial proportions of the genome (Dumont et al. 2011; Wang et al 2017) and in total number of crossovers (Dumont and Payseur 2011; Peterson and Payseur 2021), both within and between subspecies. There is also evidence that *PRDM9*, the gene that encodes the protein that dictates the locations of recombination events, has undergone recurrent bouts of positive selection in mice (Oliver et al. 2009) and natural populations of *M. musculus spp.* possess various *PRDM9* alleles corresponding to different suites of recombination hotspots (Smagulova et al. 2016). Overall, there is clear evidence from mice that recombination rates can evolve on broad and fine scales.

Changes in the recombination rate over time may influence patterns of genetic variation across the genome (Comeron 2017). For example, chromosomal fusions would decrease recombination rates experienced by individual nucleotides in the fused chromosomes, and thus increase the effects of BGS and other processes mediated by recombination. Consistent with this, Cicconardi et al. (2021) found evidence suggesting that chromosomes that underwent fusions in the ancestors of extant *Heliconius* butterfly species now exhibit reduced recombination rates and π presumably due to amplified BGS effects. Following evolution of the recombination rate landscape there will be a lag period wherein patterns of genetic variation more closely reflect ancestral recombination rates than derived rates. Over time, as new deleterious mutations arise and cause BGS, patterns of genetic variation will come to reflect derived recombination rates. Depending on the extent and rate of recombination rate evolution, population genomic analysis of lineages that are still within the lag period may be obscured. In this paper, we examine how patterns of neutral genetic variation under BGS respond to evolution of the recombination rate and describe how this could affect and have affected analyses that are used to identify the effects of selection on a genome-wide scale.

## Results

### Background selection under evolving recombination rates

The effects of BGS reflect the interplay of purifying selection and recombination (Nordborg et al. 1996), so changes to the recombination rate will influence the effects of BGS. An increase in the recombination rate between neutral sites and sites subject to purifying selection will decrease the effect of BGS and *vice versa* for a decrease in the recombination rate. At a neutral locus *v*, coalescence times under BGS (*T*_*BGS,v*_) are shorter than those expected under neutrality (*T*_*Neutral*_)(Nordborg et al. 1996) and the effect of BGS is often expressed as *B*_*v*_ = *T*_*BGS,v*_/*T*_*Neutral*_ (e.g. Nordborg et al. 1996). Consider a population that underwent a change in the recombination rate such that *v* experiences a BGS effect of *B*_*v*_′ under the derived recombination rate regime. Even with instantaneous changes in the recombination rate, genetic variation at *v* would not reflect *B*_*v*_′ immediately, as there would be a lag period after recombination rate change wherein coalescence times (and patterns of genetic variation) would more closely reflect *B*_*v*_.

Under strong purifying selection, BGS resembles a localised reduction in the effective population size, so the period of lag after a change in the recombination rate may resemble the change in coalescence times following a change in the population size. If the recombination rate changed at time *t* in the past (measured in 2*N*_*e*_ generations), then BGS under the new recombination rate can described with:

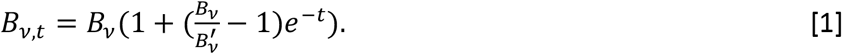

We obtained Equation 1 by modifying an expression that describes coalescence times after an instantaneous change in the population size from Johri et al. (2020). Note that Pool and Nielsen (2009) provided similar expressions to those given by Johri et al. (2020).

We modelled deleterious mutations occurring in a single functional element (e.g. a protein coding exon) and examined π for neutral mutations in and around this region after an instantaeous change in the recombination rate (Figure S1). π gradually departs from the expectations based on the ancestral recombination rate over 4Ne generations, when it finally aligns to the derived recombination rate (Figure 1,S2). Up to ~*2N*_*e*_ generations after a change in the recombination rate, π more closely resembled the expectation under the ancestral recombination rate than it did the derived rate (Figure 1, S2). After around 4*N*_'_ generations, coalescence times closely reflected those expected under BGS given the derived recombination rate, as measured by π (Figure 1, S2). When deleterious mutations have nearly neutral deleterious effects, Equation 1 may not predict changes in nucleotide diversity particularly well because in such cases BGS does not resemble a simple reduction in *N*_*e*_ (Good et al. 2014; Cvijović et al. 2018).

**Figure 1.**
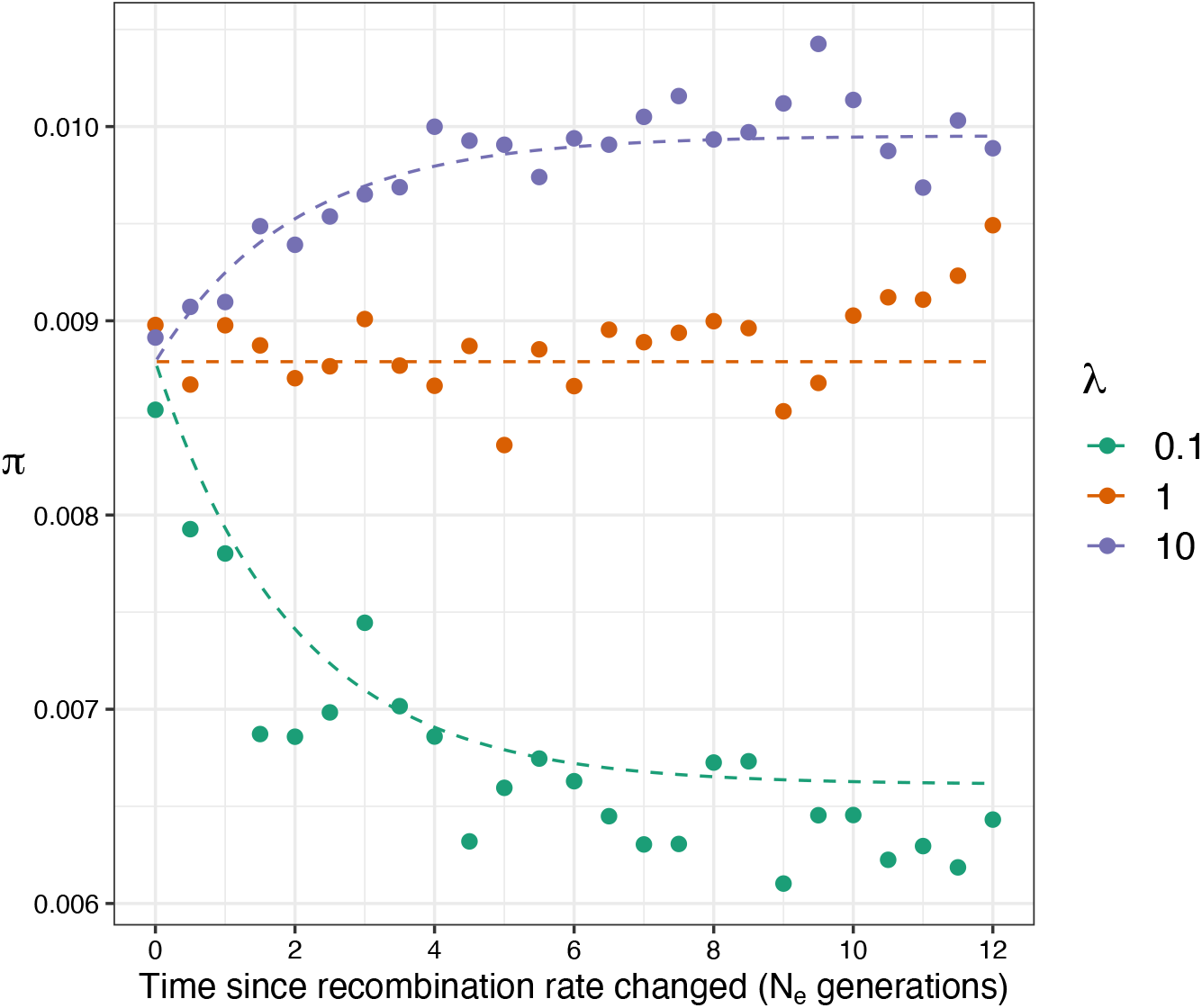
The effect of background selection on nucleotide diversity (π) over time after recombination rates change by a factor *λ*. The dashed lines were calculated using Equation 1 and formulae from Nordborg et al (1996). Points indicate the mean from 100 replicate simulations. Nucleotide diversity was calculated for neutral sites 10,000bp away from sites subject to purifying selection.

In the case of a population that has recently undergone shifts in the recombination rate landscape (i.e. less than 2*N*_*e*_ generations ago), estimates of *r* from such a population would likely reflect contemporary recombination rates regardless of how they were obtained. Estimates of *r* from patterns of marker inheritance in crosses or pedigrees always reflect contemporary rates and population genetic estimates (i.e. obtained from patterns of LD) can reflect contemporary recombination rates within 0.5*N*_*e*_ generations of a change in *r* (Figure S3). Depending on the extent and nature of recombination rate evolution, population genomic analyses that compare features of genetic variation to estimates of *r* could lead to an underestimation of the effects of BGS (and other forms of selection) on patterns of genetic variation.

### Patterns of genetic variation after evolution of the recombination landscape

To demonstrate how population genomic analyses may be affected by changes in *r*, we simulated two scenarios of BGS under evolving recombination rates. In the first, the broadscale landscape of *r* was rearranged (Figure S4A). In the second, the locations of recombination hotspots were shifted, as if a new PRDM9 allele had fixed in a population (Figure S4B). In both scenarios, deleterious mutations occurred at random across the genome generating widespread BGS such that there was a positive correlation between π and *r* at equilibrium (Figure 2). For the sake of our analyses have assumed that recombination rate is invariant among individuals, even as heritable variation in recombination rates has been reported in several species (reviewed in Stapley et al. 2017).

**Figure 2.**
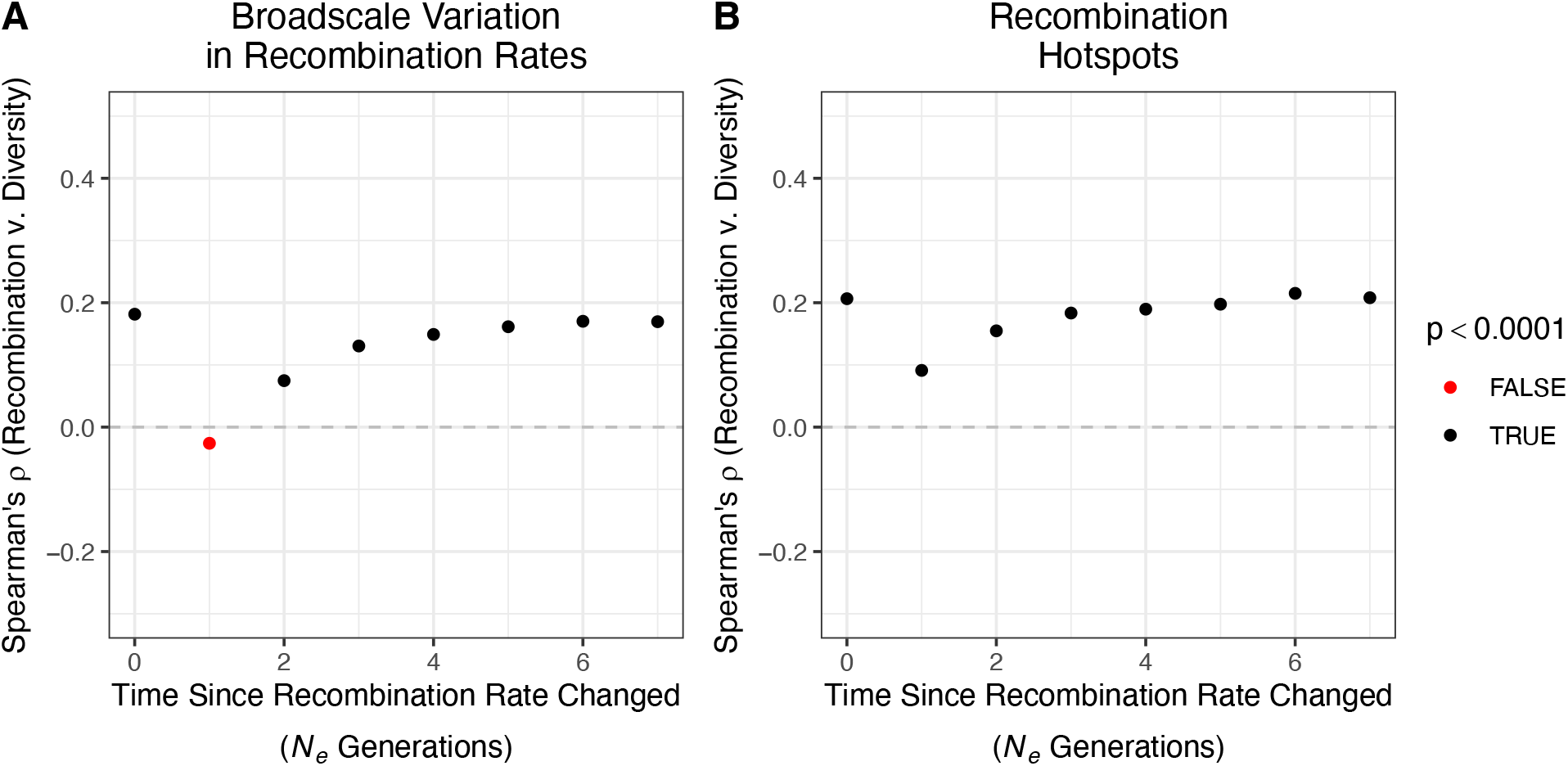
Spearman’s correlation between nucleotide diversity (π) and recombination rate (*r*) over time after recombination rates evolve. Panel A shows results for a broad-scale shift in the recombination landscape and panel B shows results for recombination rate evolution by the movement of hotspots. Results are shown for 10 Kbp analysis windows.

A positive correlation between π and *r* is a hallmark of widespread selection across a genome (Cutter and Payseur 2013), but evolution of the recombination rate may obscure this pattern. In both the scenarios we simulated, changes in *r* did not influence the average nucleotide diversity across simulated chromosomes (Figure S5), because under the models of recombination rate evolution we implemented the average map length was constant over time. However, before the change in the recombination rate, there was a positive correlation between π and *r* in both scenarios that was detectable when examining 10Kbp, 100Kbp and 1Mbp analysis windows (Figure 2,S6). Following changes in the recombination rate landscape under the model of broadscale recombination rate variation, the correlation between *π* and *r* was either absent or misleading (Figure 2A, S6A). Under the model of recombination hotspot evolution, the correlation between π and *r* was weakened by change in the landscape of hotspots (Figure 2B). In both cases we simulated, a positive correlation between π and derived *r* was restored to levels similar to what had been observed before the recombination maps changed after about 4*N*_*e*_ generations (Figure 2, S6). Figure 2 shows results for 10,000bp analysis windows, but similar results were found when examining larger windows (Figure S5).

### Rapid recombination rate evolution in house mice

Rapid evolution of recombination rates in *Mus musculus* may have influenced our ability to identify the effects of selection across that species’ genome. Kartje et al. (2020) recently demonstrated that natural populations of *M. m. domesticus* exhibit a very weak correlation between π and *r* (when examining analysis windows of various widths) and concluded that selection at linked sites exerted only modest effects on genetic variation throughout the genome. This is notable because wild mice are thought to have large effective population sizes for mammals (Leffler et al. 2012) and genome-wide effect of selection is thought to be more pronounced in species with large *N*_*e*_ (Cutter and Payseur 2013). As discussed in the Introduction, there is evidence that mice have undergone rapid evolution of the recombination rate. For example, around 3-5 MYA the lineage leading to *M. musculus* experienced a burst of karyotype evolution (Thybert et al. 2018). If that burst of karyotype evolution affected recombination rates and ancestral mouse populations were very large, then contemporary mice may still be within the lag period described by Equation 1. Patterns of genetic diversity in mice may still be adjusting to historical changes in the recombination rate, and we may see a stronger correlation between *π* and *r* in genomic regions that have not undergone dramatic changes in the recombination rate.

Using an alignment of genomes from closely related species, Thybert et al. (2018) distinguished chromosomes in the *M. musculus* genome that have or have not undergone dramatic rearrangements in the last 5 million years from those that have not. We re-analysed data from Kartje et al. (2020) and found that the correlation between π and *r* is stronger and more significant on chromosomes that have not undergone largescale rearrangements in the last 3-5 million years (Table 1) for *M. m. domesticus* individuals from France and Germany. This pattern holds when looking at analysis windows of 5 Kbp and 1 Mbp (Table 1). No substantial correlations were found for mice from Gough Island in any comparison. *M. m. domesticus* are believed to have colonised Gough Island in the 19th century and to have experienced a severe population bottleneck (Gray et al. 2014), a demographic history that could have further obscured the correlation between nucleotide diversity and recombination rate in that population.

**Table 1.** The correlation between nucleotide diversity (*π*) and recombination rate (*r*) for three populations of house mice (*Mus musculus domesticus*) calculated from all autosomes, conserved chromosomes that exhibit no syntenic breaks between *M. musculus* and *M. pahari* and chromosomes that experienced large scale rearrangements as identified by Thybert et al. (2018). Correlations with *p-*values less than 0.01 are highlighted in bold text.

| Window Size | Population | Whole Genome |  | Conserved Chromosomes |  | Rearranged Chromosomes |  |
| --- | --- | --- | --- | --- | --- | --- | --- |
| | | Spearman's $\rho$ | $p$-value | Spearman's $\rho$ | $p$-value | Spearman's $\rho$ | $p$-value |
| 5Kbp | Gough Island | **0.007 67** | **$4.28 \times 10^{-5}$** | 0.008 80 | 0.0102 | 0.004 86 | 0.0302 |
| 5Kbp | France | 0.004 08 | 0.0295 | **0.0403** | **$6.10 \times 10^{-32}$** | **-0.0107** | **$1.76 \times 10^{-6}$** |
| 5Kbp | Germany | **0.007 52** | **$6.05 \times 10^{-5}$** | **0.0152** | **$9.63 \times 10^{-6}$** | 0.003 86 | 0.0849 |
| 1Mbp | Gough Island | **0.0536** | **0.009 46** | 0.0588 | 0.124 | 0.0437 | 0.0748 |
| 1Mbp | France | 0.0450 | 0.0294 | **0.135** | **0.000 400** | 0.009 99 | 0.684 |
| 1Mbp | Germany | **0.0535** | **0.009 53** | 0.0775 | 0.0428 | 0.0426 | 0.0828 |

## Discussion

Evolution of the recombination rate will influence the effects of selection at linked sites (e.g. BGS and selective sweeps) and thus influence patterns of genetic variability. Estimates of the recombination rate made from contemporary populations may not adequately predict genetic variability up to 2*N*_*e*_ generations following evolution of the recombination rate landscape (Figure 1, 2). Our re-analysis of the Kartje et al. (2020) data suggests that mice are still within the lag period after evolution of the recombination rate, such that π in *M. m. domesticus* does not fully reflect contemporary recombination rates in *Mus musculus*. In contrast, the ancestors of *Heliconius* butterflies also underwent large-scale karyotype evolution, but gross patterns of π versus chromosome length in those species suggest that patterns of variation have largely re-equilibrated after changes in *r* (Cicconardi et al. 2021).

While our re-analysis of the data from Kartje et al. (2020) suggests that recombination rate evolution in the ancestors of mice obscures the evidence for natural selection across the genome, the overall correlations between *π* and *r* were still fairly weak on the conserved chromosomes (Table 1). The largest rank correlation coefficient we found was 0.135 for the sample of *M. m. domesticus* from France (1Mbp windows; Table 1). By contrast, Spearman’s rank correlation between nucleotide diversity and recombination rate in humans has been reported to be 0.219 for 400 Kbp analysis windows (Cai et al. 2009). The variance in recombination rates across the *M. musculus* genome is less than a half that which has been reported for humans (Jensen-Seaman et al. 2004), so perhaps the effects of BGS across the genome are more homogenous in *M. musculus* than they are in humans, contributing to the weak correlations between π and *r* shown in Table 1. Beyond the pulse of karyotype evolution reported by Thybert et al. (2018), there is clear evidence of recent and likely ongoing evolution of the recombination rate in *M. musculus* (see Introduction), which may further obscure genome-wide evidence for the effects of natural selection. For example, there is strong evidence that the landscape of recombination hotspots in the *M. musculus* genome has evolved rapidly among sub-species and populations (Smagulova et al. 2015). Our simulations suggest that even a single change to the locations of hotspots can substantially weaken the correlation between *π* and *r* (Figure 2, S6). Of course, there are reasons why species may not exhibit a strong positive correlation between π and *r* that have nothing to do with recombination rate evolution (Cutter and Payseur 2013). For example, wild and domesticated rice (*Oryza spp.*) exhibit negative correlations between π and *r*, but in those species there is a strong positive correlation between the density of functional sites (i.e. sites subject to purifying selection) and the recombination rate (Flowers et al. 2011). In such a case, the effects of BGS are primarily occurring in regions of high recombination.

This short paper should add to the growing appreciation of recombination as an evolutionarily labile trait. As pointed out by Comeron (2017) and Smukowski Heil et al. (2015), information on recombination rates in outgroup species is an important covariate when performing population genomic analyses. In some lineages, recombination rates may evolve very slowly. Birds, for example, have highly conserved karyotypes and in some cases highly conserved recombination landscapes (Damas et al. 2018; Singhal et al. 2015). Evolution of the recombination rate is another of the many possible reasons why one might not be able to adequately identify the effects of BGS (or natural selection more broadly) from population genomic data (See reviews by Cutter and Payseur 2013 and Comeron 2017), but conservation of recombination landscapes will likely make comparative population genomics more straightforward.

## Methods

### Model

Background selection has been modelled as the reduction in effective population size (*N*_*e*_) at a neutral site due to the removal of linked deleterious variants. The effects of background selection are often expressed as 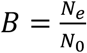, where *N*_*e*_ is the effective population size and *N*_0_ is the expected population size under strict neutrality. In a non-recombining genome, *B* is proportional to the ratio of the deleterious mutation rate to the strength of selection acting on harmful mutations (Charlesworth et al. 1993). For a neutral site present on a recombining chromosome, the effects of background selection depend on the density of functional sites (i.e. those that can mutate to deleterious alleles), the strength of selection at functional sites, the mutation rate at functional sites and the recombination rate between the neutral site and the functional sites (Hudson and Kaplan 1995; Nordborg et al. 1996; Nordborg 1997). For a neutral locus *v* linked to *x* functional sites, the reduction in *N*_*e*_ has been described with the following equation:

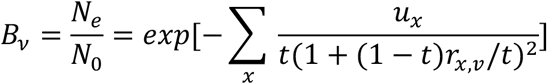

where *u*_*x*_ is the deleterious mutation rate at functional site *x*, *t* is the heterozygous fitness effect of a deleterious mutation (i.e. 0.5*s* in the case of semi-dominance) and *r*_*x,v*_ is the recombination map distance between the neutral locus and functional site *x*. In the above equation, deleterious mutations have fixed effects, but it is straightforward to incorporate a distribution of fitness effects (Nordborg et al. 1996). The above equation holds when selection is sufficiently strong such that random drift does not overwhelm selection (*N*_*e*_*s* > 1) (Good et al. 2014).

### Simulations

We simulated BGS under recombination rate evolution using two types of simulations in *SLiM* v3.2 (Haller and Messer 2019). We simulated diploid populations of *N*_*e*_ = 5,000 individuals. In all cases, we scaled mutation, recombination and the strength of selection to approximate evolution in a large population.

The first set of simulations was designed to examine how long it takes for patterns of neutral diversity under BGS to equilibrate after the recombination rate evolves. In these simulations, the genome was 25 Kbp long with a 5 Kbp functional element in the centre. Mutations occurred in the functional element at rate *μ* = 5 × 10^−7^ and had semi-dominant fitness effects with a fixed selection coefficient of *s* = −0.01. We also simulated cases with varying fitness effects using a gamma distribution with mean 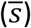 of −0.1 and a shape parameter of 0.1. Recombination occurred at a uniform rate of *r* = 5 × 10^−7^ across the chromosome. After 80,000 generations (16*N*_*e*_ generations), we simulated an instantaneous change in the recombination rate, multiplying *r* by *λ*, giving *r* = *λ*5 × 10^−7^. We simulated cases with *λ* = 0.1, 1.0 and 10.0. Simulated populations were sampled every 2,500 generations after the recombination rate changed and we performed 200 replicates for each set of parameters tested. Note that these simulations were not designed to be particularly realistic, but to provide clear cut patterns to test the theoretical predictions.

The second set of simulations was designed to examine how patterns of π versus *r* varied over time when recombination rates evolved at fine and/or broad scales. For these simulations, we modelled chromosomes that were 10 Mbp long. Neutral mutations occurred at random across the length of the sequence at a rate of 5 × 10^−7^ (such that expected nucleotide diversity was 0.01). Deleterious mutations occurred at random across the length of the sequence at a rate of 5 × 10^−8^ with semi-dominant fitness effects drawn from a gamma distribution with a mean 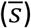 of −0.1 and a shape parameter of 0.1. The deleterious mutation rate was chosen so that 10% of the genome was subject to purifying selection. Populations evolved under background selection for 80,000 generations (i.e. 16*N*_*e*_ generation). In generation 80,000 there was instantaneous evolution of the recombination landscape after which we recorded the tree-sequence of the population every 5,000 generations for a further 40,000 generations. We incorporated two models of recombination rate variation and evolution of the recombination map:

- We modelled recombination rate evolution at broad scales by rearrangement of the recombination landscape. Recombination rates vary across the genome (Stapley et al. 2017). For example, recombination rates vary by a factor of 3 across chromosome 1 in mice. In these simulations, recombination varied from *r* = 2.08 × 10^−7^ to *r* = 6.24 × 10^−7^ across the simulated chromosome (Figure S4A). When the recombination landscape evolved, we reversed the order of recombination rates across the genome (Figure S4A).
- We modelled evolution of the recombination map by the movement of hotspots. Recombination occurred at a uniform rate of *r* = 6 × 10^−8^ except in 5 Kbp hotspots where it occurred at a rate of *r* = 6 × 10^−6^. At the beginning of a simulation, a Poisson number of hotspots was sampled with an expectation of 120. Hotspots were placed at random across the simulated chromosome. When the recombination landscape evolved, we resampled the locations of hotspots (Figure S4B).

In both cases, rates were chosen such that the total map length was similar to one that recombined at a constant rate of 4*N*_*e*_*r* = 0.008, the value reported for wild mice (Booker et al. 2017). For both models of recombination rate map evolution, we performed 20 simulation replicates, giving a total of 200 Mbp worth of simulated data, similar to the length of chromosome 1 in mice.

For all simulations, we used the tree sequence recording option in *SLiM* and neutral mutations were added to the resulting tree-sequences at a rate of 5 × 10^−7^ using *PySLiM* and *msprime* (Haller et al. 2019; Kelleher et al. 2016). Nucleotide diversity (π) was calculated in windows of varying size using sci-kit-allel. We used the program *PyRho* (Spence and Song 2019) to estimate recombination rates from samples of 10 diploid individuals from 20 replicate simulations. Spearman’s *ρ* between π and *r* was calculated using R. All figures were made using ggplot2. All simulation scripts and analysis and plotting scripts are deposited at https://github.com/TBooker/BGS_RecombinationRateEvolution.

## Acknowledgements

Thanks to Nadia Singh and Judith Mank for the invitation to present this work at vSMBE 2021. Thanks to Michael Whitlock, Brian Charlesworth and Mikey Kartje for helpful discussions. TRB was funded by a Bioinformatics Fellowship awarded by the Biodiversity Research Centre at the University of British Columbia. BAP was supported by NIH R35 GM139412.

## Supplementary Material

**Figure S1.**
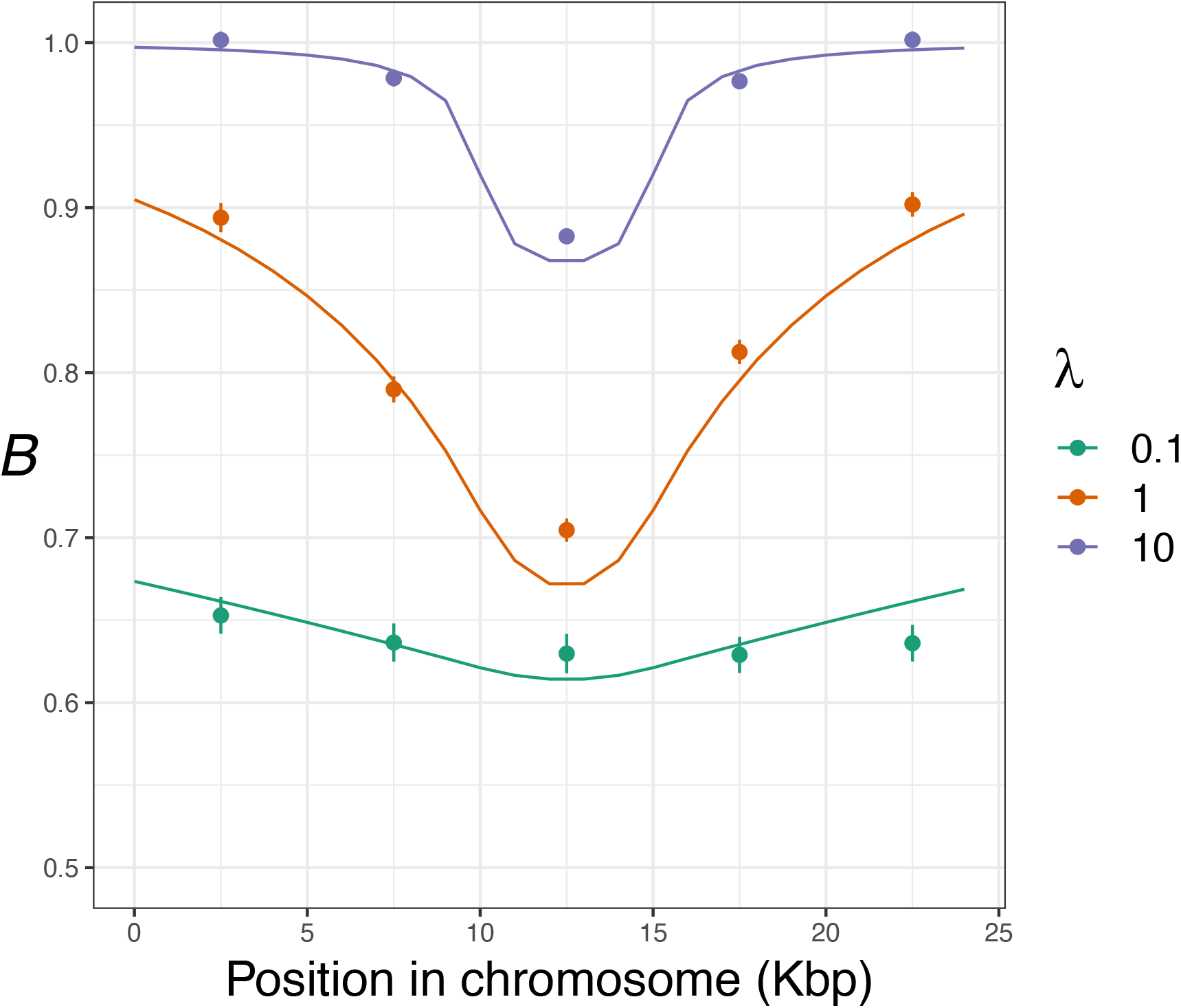
The effects of background selection across simulated chromosomes. B was calculated for simulated data by comparing observed π to the neutral expectation of 4*N*_*e*_*μ* = 0.01. The lines show the theoretical expectation calculated using formulae from Nordborg et al (1996).

**Figure S2.**
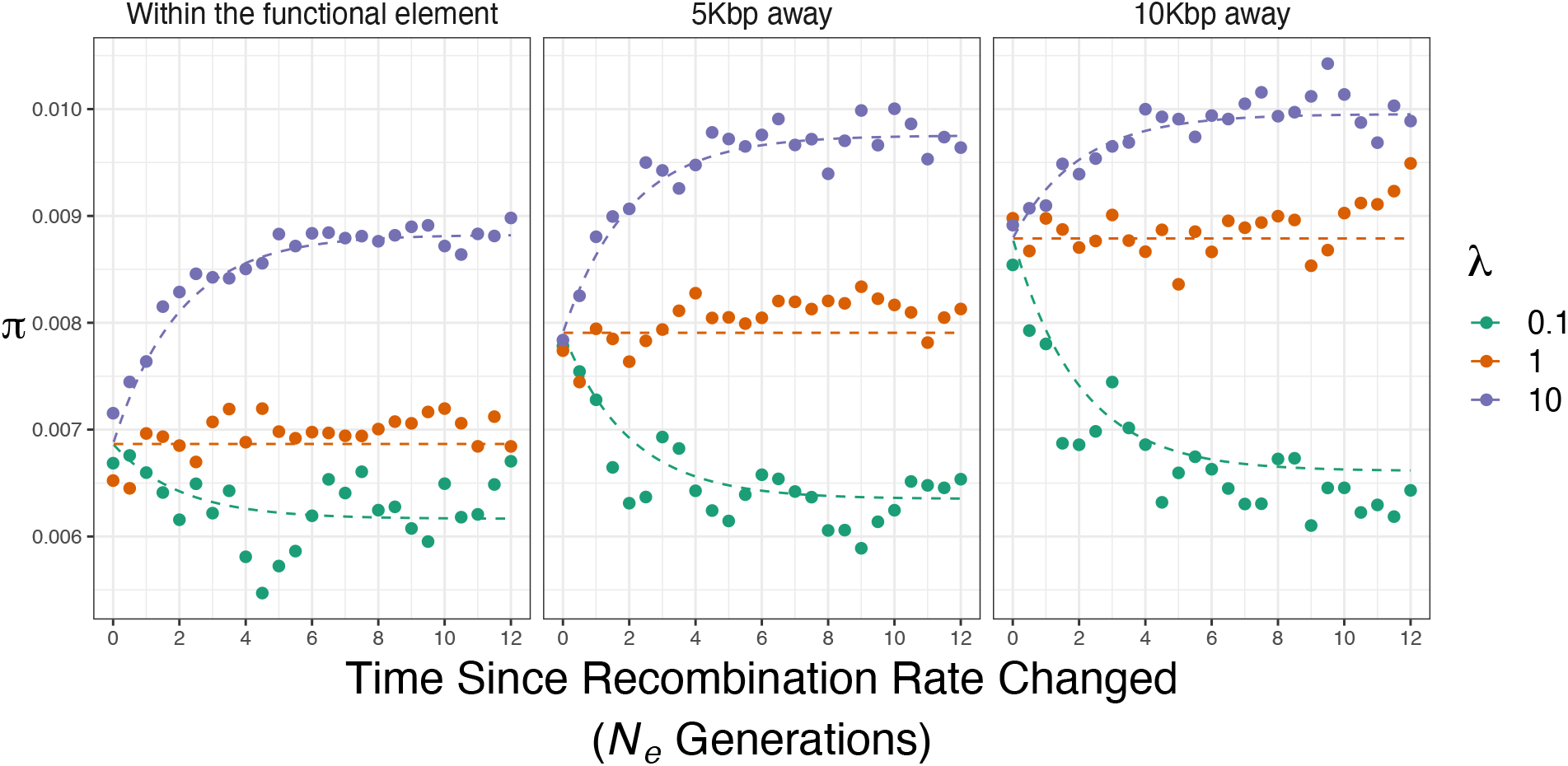
The effects of background selection across simulated chromosomes. B was calculated for simulated data by comparing observed π to the neutral expectation of 4*N*_*e*_*μ* = 0.01. The lines show the theoretical expectation calculated using Equation 1 and formulae from Nordborg et al (1996). The labels on the top of each panel indicate the location in the simulated data being analysed.

**Figure S3.**
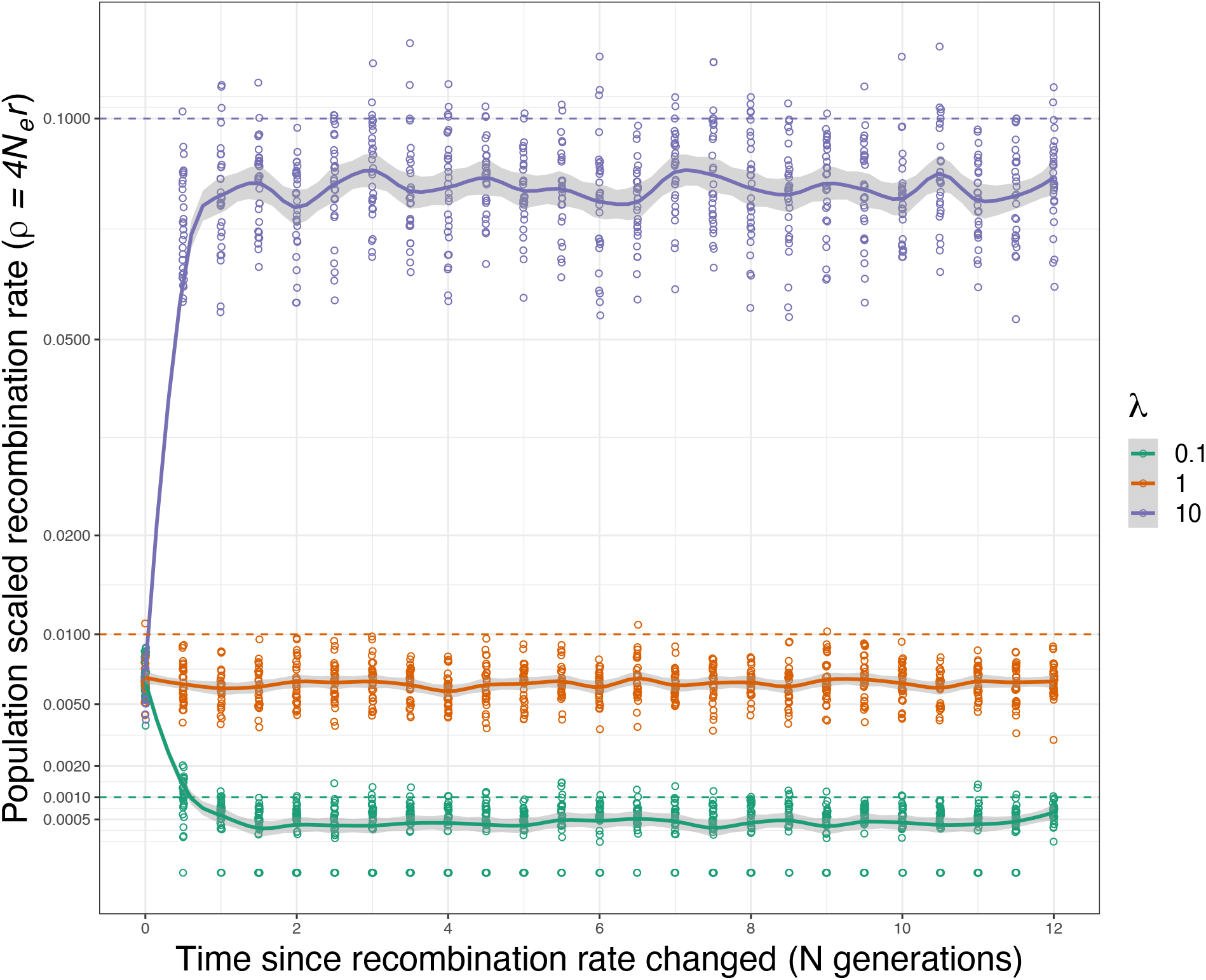
Recombination rates inferred using PyRho after an instantaneous change in the recombination rate. Dashed horizontal lines indicate the true recombination rate for the three cases. Smoothed lines with shaded ribbon indicate the fit and error of a LOESS regression. Recombination rates were estimated for 30 simulation replicates for each time point and value of λ.

**Figure S4.**
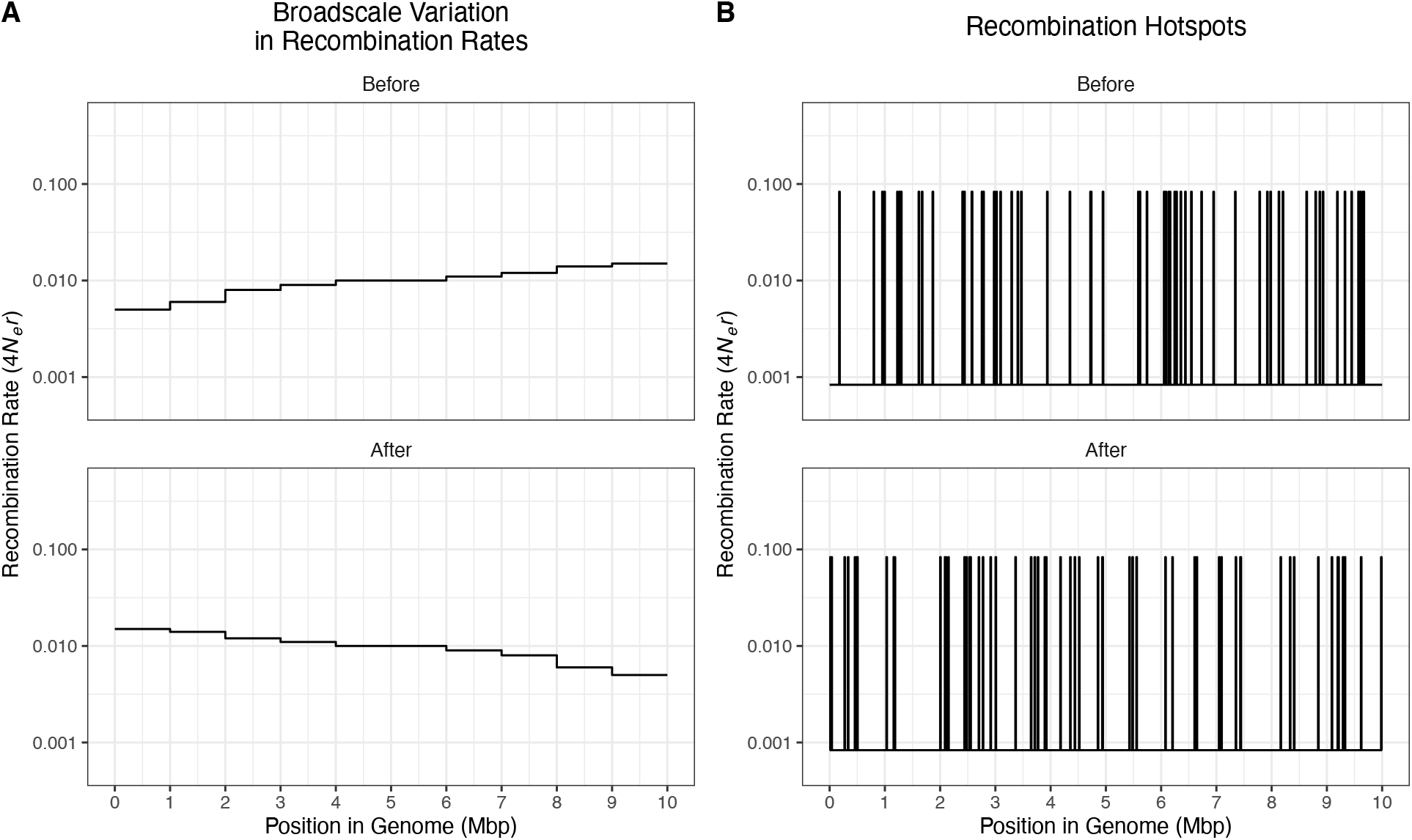
The recombination rate maps used in the simulations modelling BGS across the genome. The upper and lower panels show the recombination rate landscape before and after it evolved in simulations, respectively. A) Evolution of the recombination rate at the Mbp scale. B) Evolution of the recombination rate at the scale of recombination hotspots.

**Figure S5.**
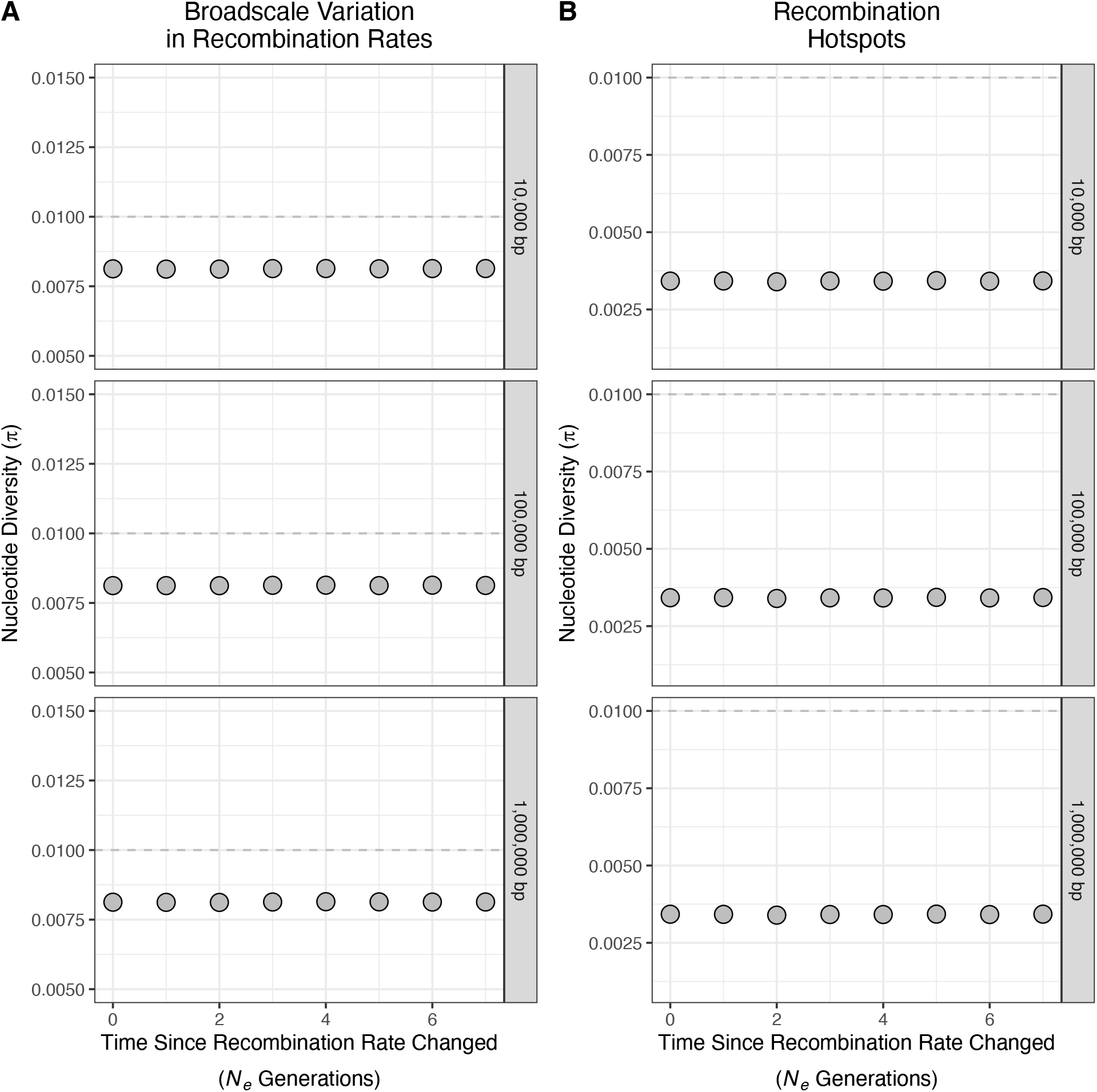
Nucleotide diversity (π) over time after evolution of the recombination landscape. Panel A) shows results for the model of broadscale recombination rate evolution. Panel B) shows the results for the model of recombination hotspot evolution. The dashed horizontal grey line indicates the null expectation of *4N*_*e*_*μ* = 0.01. In both panels, the text in the grey strips to the right of each cell indicates the size of analysis windows used.

**Figure S6.**
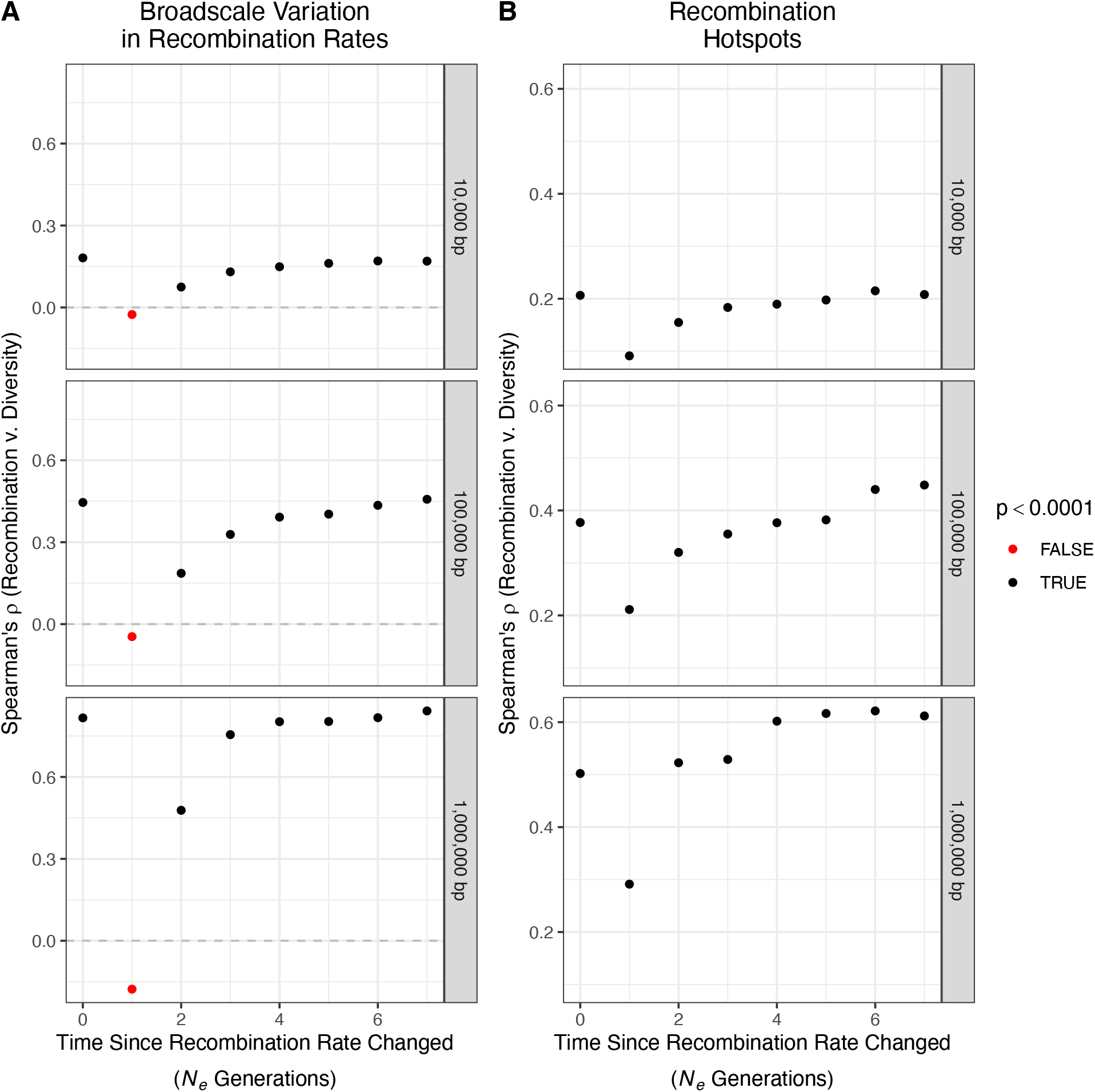
The correlation between nucleotide diversity and recombination rate over time after evolution of the recombination landscape. Panel A) shows results for the model of broadscale recombination rate evolution. Panel B) shows the results for the model of recombination hotspot evolution. In both panels, the text in the grey strips to the right of each cell indicates the size of analysis windows used.

